# EdiTyper: a high-throughput tool for analysis of targeted sequencing data from genome editing experiments

**DOI:** 10.1101/2020.07.30.229088

**Authors:** Alexandre Yahi, Paul Hoffman, Margot Brandt, Pejman Mohammadi, Nicholas P. Tatonetti, Tuuli Lappalainen

## Abstract

Genome editing experiments are generating an increasing amount of targeted sequencing data with specific mutational patterns indicating the success of the experiments and genotypes of clonal cell lines. We present EdiTyper, a high-throughput command line tool specifically designed for analysis of sequencing data from polyclonal and monoclonal cell populations from CRISPR gene editing. It requires simple inputs of sequencing data and reference sequences, and provides comprehensive outputs including summary statistics, plots, and SAM/BAM alignments. Analysis of simulated data showed that EdiTyper is highly accurate for detection of both single nucleotide mutations and indels, robust to sequencing errors, as well as fast and scalable to large experimental batches. EdiTyper is available in github (https://github.com/LappalainenLab/edityper) under the MIT license.

## 1 Introduction

Genome editing techniques have quickly become an essential tool for molecular biology, with diverse applications in many fields. This revolution has been driven in particular by the effi cient CRISPR-based genome editing [7, 3, 1, 5], based on a guide RNA (sgRNA) that can target a Cas9 or other nuclease to a specific sequence to introduce a double stranded break (DSB) of DNA. Its repair by non-homologous end-joining (NHEJ) or homology-directed recombination (HDR) can be used to induce small indels or specific edits, often single base edits [13].

In applications that aim to edit a specific set of loci, targeted sequencing on the Illumina platform is the most common approach to determine the outcome of a genome editing experiment. However, analysis of the resulting sequencing data is not always straightforward. Repair of double-stranded breaks introduces diverse types of mutations, and especially the analysis of polyclonal populations of edited cells contain a diverse set of mutations that is not well amenable to standard genotyping tools. Furthermore, the ideal tool would be not only accurate but also fast enough to enable analysis of large amounts of sequencing data from thousands of monoclonal cell lines, be easy to install and use even by non-experts, and provide comprehensive output data for downstream analysis and troubleshooting.

A number of tools have been developed for analysis of sequencing data from CRISPR experiments. Fully web-based tools such as CRISPR-GA [4] are not scalable to very large experiments. More recent tools include AGEseq [14] with a standalone program along with a Galaxy-based web tool, but an output that only offers a table summary. The open source CRISPResso [11] offers both command-line and web-based tools, with features such as batch processing, figures and detailed outputs. Cas-analyzer [10] presents an hybrid solution by offering a web-based tool on the client side, virtually suppressing the need of uploading data on distant servers, therefore getting rid of associated security risks, and offering a graphic interface instead of a command-line tool. While CRISPR-GA and AGEseq rely on BLAT aligner [8], CRISPResso and Cas-analyzer both use the EMBOSS Needle alignment software.[9] Since this study was conducted, CRISPResso has been updated to a new version, CRISPRESSO 2 [2], that has faster processing time and can be found at https://github.com/pinellolab/CRISPResso2. We did not use this new version in our comparisons.

In this paper, we present EdiTyper, an open-source command line software for characterizing targeted sequencing data from polyclonal and monoclonal cell lines from genome editing experiments. EdiTyper uses our novel Needleman-Wunsch alignment software, RecNW [15], is highly adaptable, and provides detailed output data. We show that EdiTyper outperforms previous software for similar purposes by being highly accurate, robust, and with high processing speed.

## 2 Methods

### 2.1 Overview and input and output data

EdiTyper is an open source command line python-based tool designed to process targeted DNA sequencing data of polyclonal or monoclonal CRISPR genome editing experiments, in order to characterize the type, frequency, and position of genome editing events (Figure 1). EdiTyper input and output data are designed to allow both detailed inspection of a small number of experiments as well as streamlined analysis of thousands of samples in parallel. The input of Illumina single- or paired-end sequencing data in EdiTyper is in the raw fastq format that can be compressed, and provided either as a single file or as a directory with several files. Additionally, a reference sequence that represents the locus before gene editing is required, and optionally the template for homologous recombination if that was used in the experiments. Metadata can be defined by the user. The processing steps include a quality-control and read pre-processing module, an alignment step using a fully customizable semi-global aligner implemented in C++ [15], and a classification module that analyzes the aligned sequences and quantifies the amount of HDR, NHEJ, HDR+NHEJ, and their position relative to the reference sequence. The specific parameters of the semi-global aligner are modifiable, such as the gap opening and gap extension penalties, along with the quality threshold used to filter out reads with a poor alignment score. The outputs can be suppressed, with the full set of files including read alignments in the SAM/BAM format that can be viewed in various other software, a table counting the events for each position on the reference, a classification file returning the label attributed to each input read, a summary table with the final read classification counts, a pdf file of figures of the alignment quality distribution, and a landscape of the editing events in the locus.

**Figure 1:**
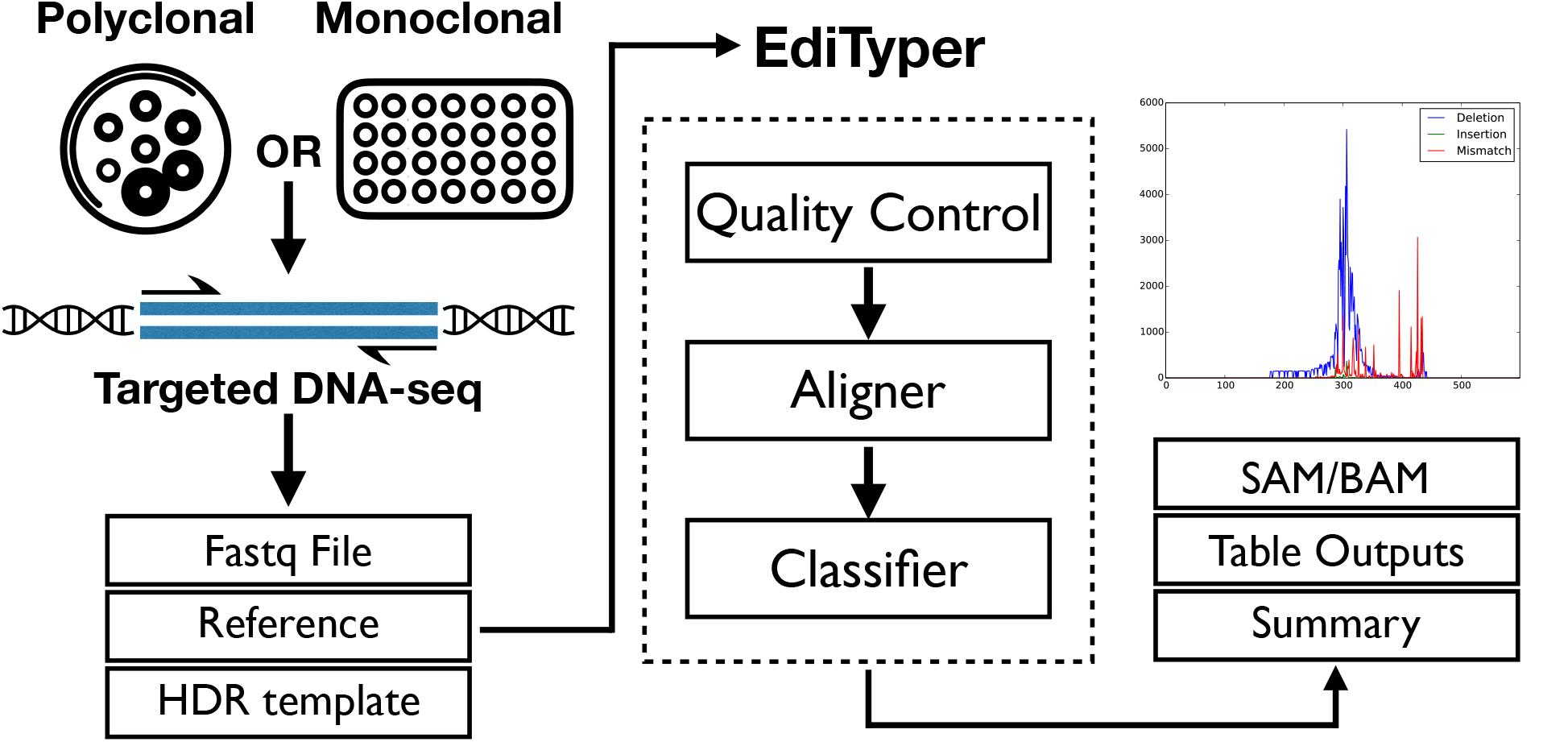
Outline of EdiTyper. It is designed for analysis of CRISPR genome editing experiments, either polyclonal cell populations containing a mixture of cells with diverse mutations, or monoclonal cells lines derived from a polyclonal population. The editing events are analyzed from a fastq file of targeted DNA sequencing on the Illumina platform. Additionally, EdiTyper requires an input of the targeted reference sequence, and optionally the HDR template to classify the edited reads. The tool itself is composed of a read preprocessing and quality control module, our custom aligner RecNW implemented in C++, and the read classification module. The EdiTyper (optional) outputs include alignments in sam/bam format, plots to describe mutations at the locus, and extensive summary files.

### 2.2 Read pre-processing and quality control

EdiTyper supports single and paired-end reads that are already trimmed for adaptor and low quality bases using standard tools. First, EdiTyper performs reverse complementation, if needed, and alignment quality filtering. This is done by taking 10% of the reads up to 500 reads, and creating 4 groups: the original reads, their reverse complement, and a permuted version of each group. All groups are aligned to the targeted reference, and alignment scores between the original and reverse complement reads are used to determine the correct orientation. The alignment scores for the permuted reads are used as a null distribution to set the alignment quality threshold, *s*^∗^, as:

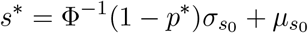

Where Φ^−1^ is the inverse of the standard normal cumulative distribution function, 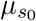 and 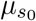 are mean and standard deviation of alignment scores from the permuted groups, and *p*^∗^ is the p-value threshold selected. This leads to inclusion of reads that align substantially better than random sequence, but the threshold can also be manually adjusted.

### 2.3 Alignment and read classification

Targeted sequencing analysis of specific loci typically produces high read counts from known loci, with high similarity between reads and an atypical mutation profile due to NHEJ, HDR, and their mixture. Thus, for alignment, EdiTyper uses a local alignment of sequencing reads to the targeted locus with RecNW, our novel software that implements a modified version of the Needleman-Wunsch algorithm [15]. RecNW provides identical results as gold-standard Needleman-Wunsch implementations in substantially higher speed for high-coverage loci, achieved by deduplication of reads and reuse of identical parts of the alignment matrices between reads.

In EdiTyper, aligned reads are passed on to a python module for classification of each read based on its mutation events (Figure 2). If a homologous template is provided, reads that carry the designated SNP mutation are considered to have HDR, and if indels are observed anywhere in the read, it is classified as NHEJ. Reads with both are classified as MIX. EdiTyper allows designation of an additional edited SNP – typically to break the PAM site – and other mismatches in the reads are reported but do not affect read classification.

**Figure 2:**
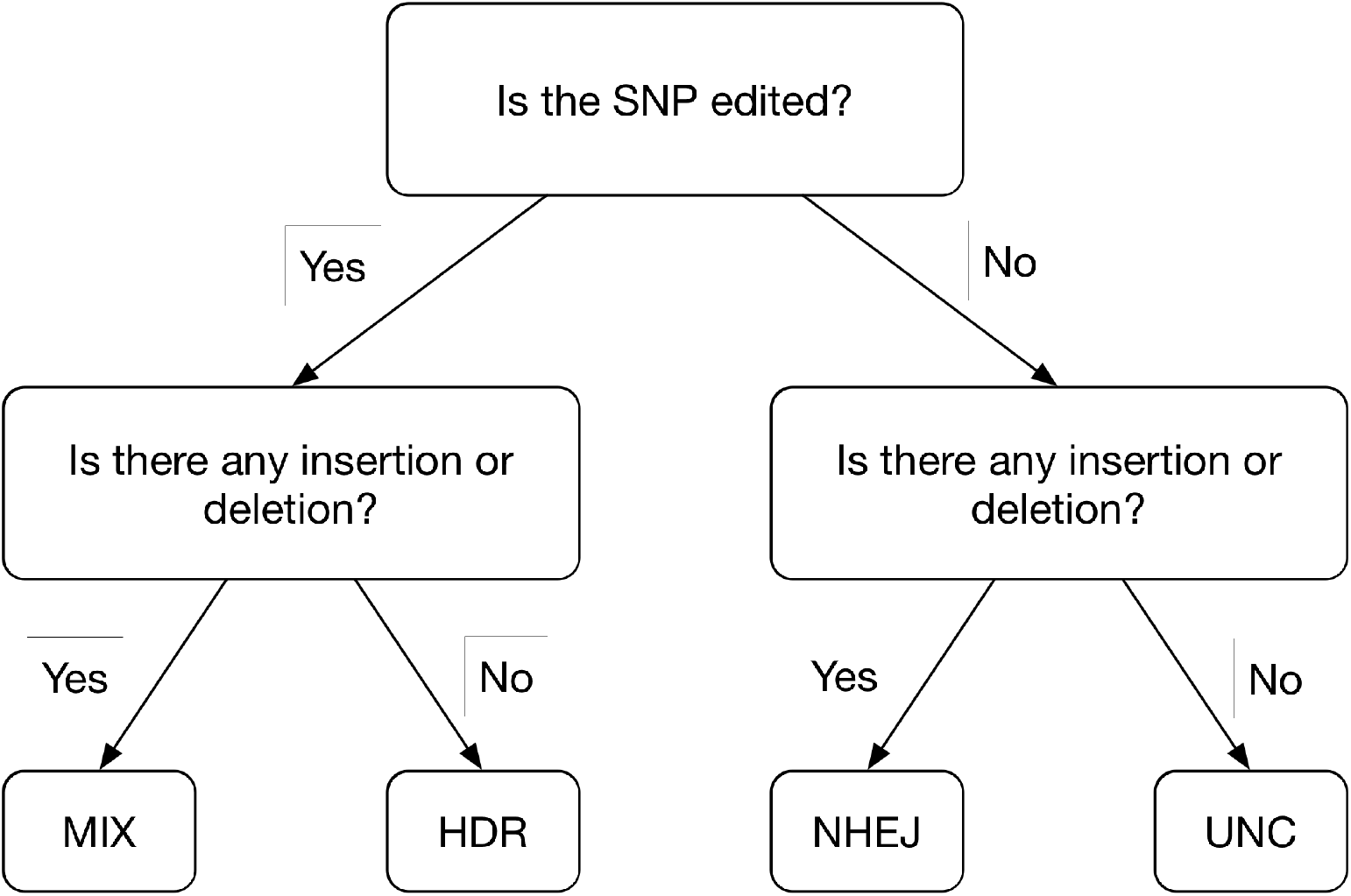
Classification decision tree to assign a label to each read based on the analysis of the alignments.

### 2.4 Implementation

EdiTyper is a software package written in Python, C++, and R. The package provides the EdiTyper command-line tool for use in pipelines, and the edityper Python module for use in custom Python scripts. Alignment is done using the C++ RecNW aligner [15], analysis is done in Python, and plotting is done in R. EdiTyper works with both Python 2, from 2.7 and up, and Python 3, from 3.5 and up, and supports both UNIX and Windows systems. Python’s multiprocessing library is used to implement parallelized processing and analysis of data. The software depends on the Cython, Biopython, and regex Python modules, as well as SAMtools for BAM file generation and R 3.1.0 or higher for plotting. The Python modules are automatically installed when installing EdiTyper with the pip package manager; SAMtools and R are user-provided and if not present, EdiTyper will skip the steps that use these tools. EdiTyper is distributed under the MIT license and is available on the Python Package Index and on GitHub.

### 2.5 Benchmarking

We first evaluated and demonstrated EdiTyper’s performance by using simulated data. We simulated reads from 17 protein-coding gene loci, with a GC% ranging from 34.67% to 82.67% (Supplementary Table 1). For each locus, we generated synthetic reads with varying parameters. We varied the percentages of HDR (0.2, 0.5, 1.0, 1.5, 2.0, 2.5, 3.0, 3.5, 4.0, 4.5 or 5%), set the percentage of MIX (HDR+indel) identical to that of HDR, and fixed the percentage of unchanged reads to 50%. The lengths of the indel for NHEJ events was random within set ranges of (1,1), (2,4), (5,10), (11, 20) bps, for NHEJ events the insertion:deletion ratio was set to 1:3, the indel position in the read was random, and the total number of reads was set to 4000, 8000, 12000, or 16000. For NHEJ and MIX reads. These reads were then processed with the ART tool to simulate sequencing error profiles from Illumina MiSeq v1 and v3 platforms [6], and we focused most analyses on these data sets since they are more realistic than reads without errors. Altogether, we generated a total of 8,976 synthetic datasets by varying the parameters in all possible combinations, or 528 datasets from each of the 17 loci.

We evaluated the classification performance and speed of EdiTyper compared to CRISPResso with the simulated data described above. CRISPResso was selected because has been is widely used, and it uses an exact alignment algorithm (i.e., EMBOSS Needle) comparable to recNW in EdiTyper, and that provides a local command line software for CRISPR experiment sequencing data analysis with similar outputs. We did not benchmark against later versions of CRISPResso and all results have been obtained using the version 1.0.X of that software https://github.com/lucapinello/CRISPResso. For the evaluation of both tools we considered processing time and classification metrics associated to the full characterization of every single read of the dataset: global accuracy, and class-specific precision and recall. We also analyzed the summary statistics of each data set to calculate the absolute and relative errors of estimation of the proportion of HDR, NHEJ, MIX and unchanged reads. We ran all experiments on a server with 64 CPU cores AMD Opteron 6272 and 322 Gb of RAM running under Ubuntu 14.04 with Python 2.7.8, g++ 4.8.4, Cython 0.24, NumPy 1.11.3 and SciPy 0.19.1.

In order to demonstrate the performance of EdiTyper on real data, we analyzed the sequencing results of an editing experiment to introduce SNP rs131811 into the 293T human cell line. After CRISPR editing with an HDR template, genomic DNA was extracted from the polyclonal cell population, and a 180 bp segment was amplified by PCR. A sequencing library was prepared from the PCR amplicon using nextera indexing, which was sequenced on the Illumina MiSeq with paired-end 150 bp reads. After adaptor and quality trimming with Trimmomatic, we ran EdiTyper on the sequencing data with default settings. The run was done on 40-core Intel Xeon 2.8 GHz CPU that has 256 GB of RAM, and runs CentOS 7.4.1708, and for estimating speed, it was run 500 times without parallelization.

## 3 Results

### 3.1 Classification accuracy

First, we compared the estimated HDR and NHEJ percentages against the true percentages used to generate our datasets, and observed a high overall concordance across the different data sets. Analyzing reads with sequencing error profiles (MSv1 and MSv3), EdiTyper made a relative error of 4.6%±6.53 (0.06%±0.047 in absolute error) on HDR estimates, and 1.5%±0.93 (0.66%±0.424 in absolute error) on NHEJ estimates (Figure 3A; see Supplementary Figures 3-4 for datasets without simulated sequencing error and results by sequencing platform). This is substantially higher than the relative errors obtained with CRISPResso of 40.2%±33.32 for HDR and 56.6%±44.65 for NHEJ (Figures S5-6). EdiTyper relative error on HDR estimates was larger than the one of NHEJ due to the low percentages of HDR in our datasets (i.e., ranging from 0.2% to 5%), but the absolute error was extremely low.

**Figure 3:**
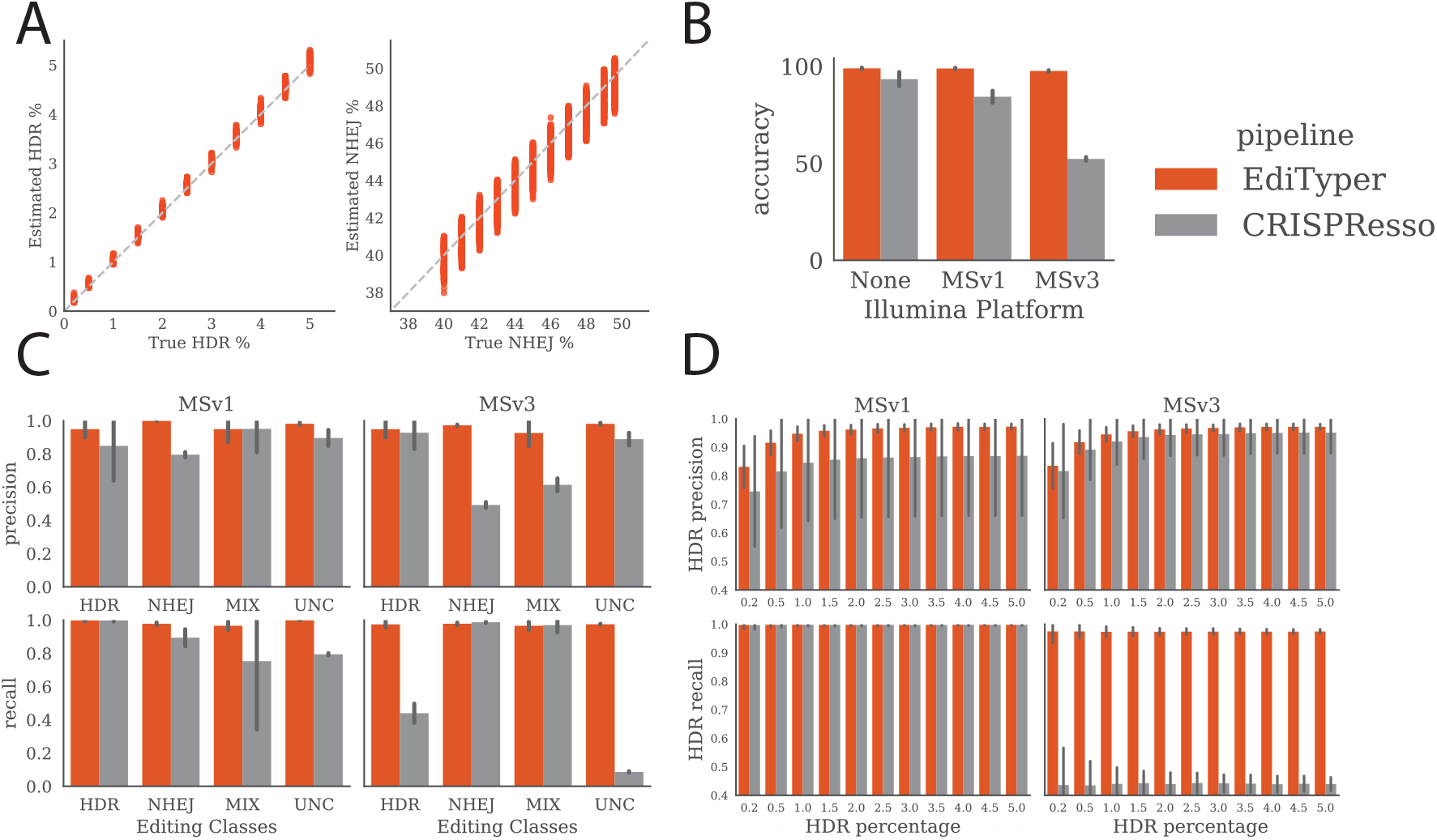
Accuracy of EdiTyper and its comparison to CRISPResso using simulated data. (A) comparison of the estimated vs. true HDR and NHEJ percentages for EdiTyper across all datasets with simulated sequencing error; (B) global accuracy between the two tools for different sequencing error modes; (C) breakdown of precision and recall for each class: homology-directed repair (HDR), non-homologous end-joining (NHEJ), MIX (HDR + NHEJ) and unchanged reads (UNC); (D) HDR precision and recall as a function of HDR percentage.

We then tested the effect of sequencing error profiles on the accuracy of both tools. Without sequencing error, both tools performed reasonably well, with EdiTyper having an average accuracy of 98.95%±0.355 and CRISPResso 93.39%±3.665. However, in the presence of sequencing error, EdiTyper was substantially more accurate (98.87%±0.353 for MSv1 and 97.59%±0.373 for MSv3) compared to CRISPResso (84.30%±3.086 fo MSv1 and 52.26%±0.947 for MSv3). These differences may be caused by MSv3 having a lower average quality than MSv1 especially in the beginning of the read, but EdiTyper has high accuracy regardless of the sequencing error profile.

EdiTyper’s accuracy was between 97.09% and 99.61% (95% interval). It was slightly affected by the average indel size (Pearson *r* = −0.19, p-value< 10*e* − 69) (Figure S7), and was negatively correlated with the GC% of the locus (Pearson *r* = −0.78, p-value= 0.0002; Figure S8). However, since the average accuracy ranges from 98.23% to 98.68% depending on the locus, the effect is negligible. To ensure good performance across all read types (HDR, NHEJ, MIX and Unchanged), we calculated precision and recall for read classification for each simulated data set, focusing on those with MSv1 and MSv3 sequencing errors. EdiTyper had a high and consistent precision and recall above 94% for all types of reads, while CRISPResso showed a mix of decreased precision or recall for various read types and sequencing error profiles, especially for MSv3 (Figure 3C; Tables S2-3). As HDR quantification is of particular importance for many genome editing experiments, we evaluated the precision and recall for HDR classification with varying percentages of HDR reads across MSv1 and MSv3 error profiles. We observed that recall was not affected by the percentage of HDR in the dataset, but HDR classification precision was decreased for both EdiTyper and CRISPResso for the very lowest percentages of HDR, likely due to reads carrying the mutation getting scarce. However, overall our simulation results indicate that EdiTyper performs very well across diverse parameters, with superior performance to an analogous previously published tool.

### 3.2 Processing speed

We measured speed both in terms of total processing time, and number of reads processed per second. Over our 8,976 simulated datasets of varying sizes, EdiTyper’s average speed was of 2255 reads/sec (455 reads/sec to 12,698 reads/sec), 12 to 18 times faster compared to CRISPResso (average of 143 reads/sec ranging from 57 reads/sec to 248 reads/sec) (Figure 4). The highly variable processing speed for EdiTyper is due to its aligner that collapses identical reads and leverages on similarities between reads (Ref: RecNW). Thus, EdiTyper has a faster processing time per read as the total number of reads increases (for example with MSv3, 532±26 reads/sec for datasets of 4000 reads versus 731±40 reads/sec for 16,000 reads), and faster processing time when there are fewer sequencing errors (average 4077 reads/sec with no sequencing errors, 2043 reads/sec for MSv1, and 645 reads/sec for MSv3). We showed that EdiTyper’s processing time is linear with the number of unique reads. Therefore it is not impacted by the total size of the dataset like CRISPResso (Figure S9).

**Figure 4:**
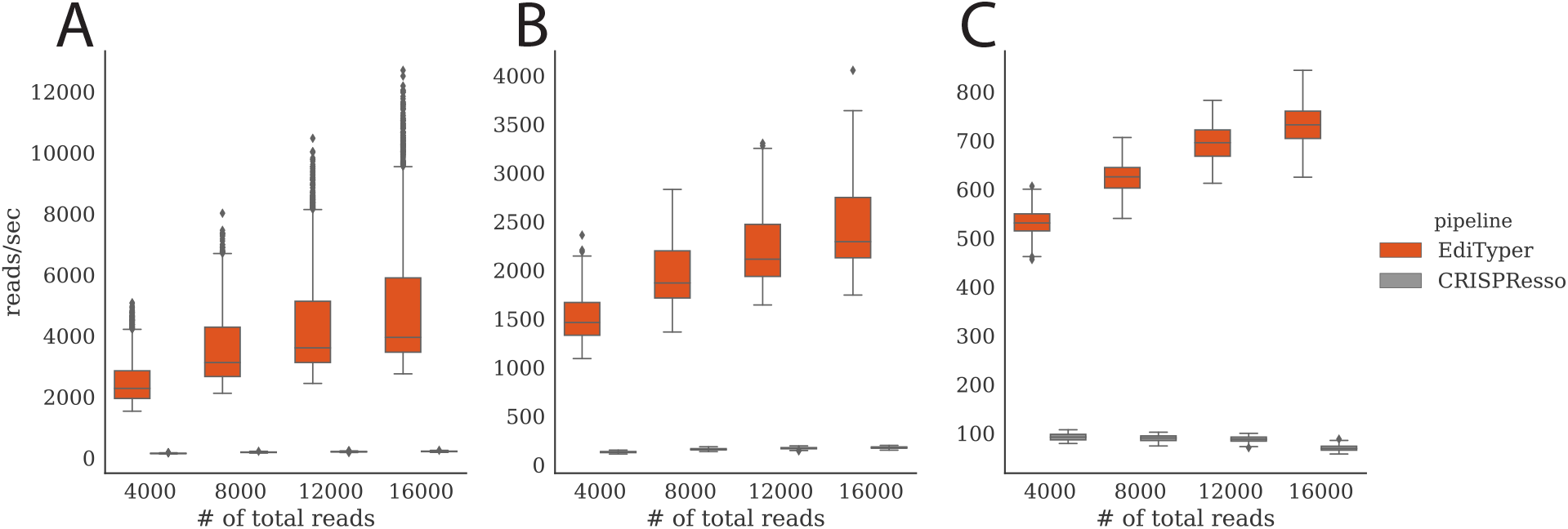
Comparison of the processing speed between EdiTyper and CRISPResso in reads/sec for data a) without sequencing error, and with b) MSv1 and c) MSv3 sequencing error profiles.

### 3.3 Results of real sequencing data

We tested EdiTyper on MiSeq sequencing data from a locus where SNP rs131811 was introduced into the human 293T cell line. A total of 70,649 reads were aligned, and 69,625 well-aligned reads were used for analysis (Figure 5): each read was classified as in Figure 2, the mutation events and coverage were recorded per base pair, and these data were written in summary files, together with alignments in bam format that can be further viewed e.g. in IGV [12], and summary plots. The analysis took a median of 88 seconds, of which 79 seconds was for the optional creation of the sam file and its conversion to bam. This demonstrates the speed of EdiTyper even for large numbers of reads from targeted sequencing. Figure 5 shows an example of two summary plots of the results.

**Figure 5:**
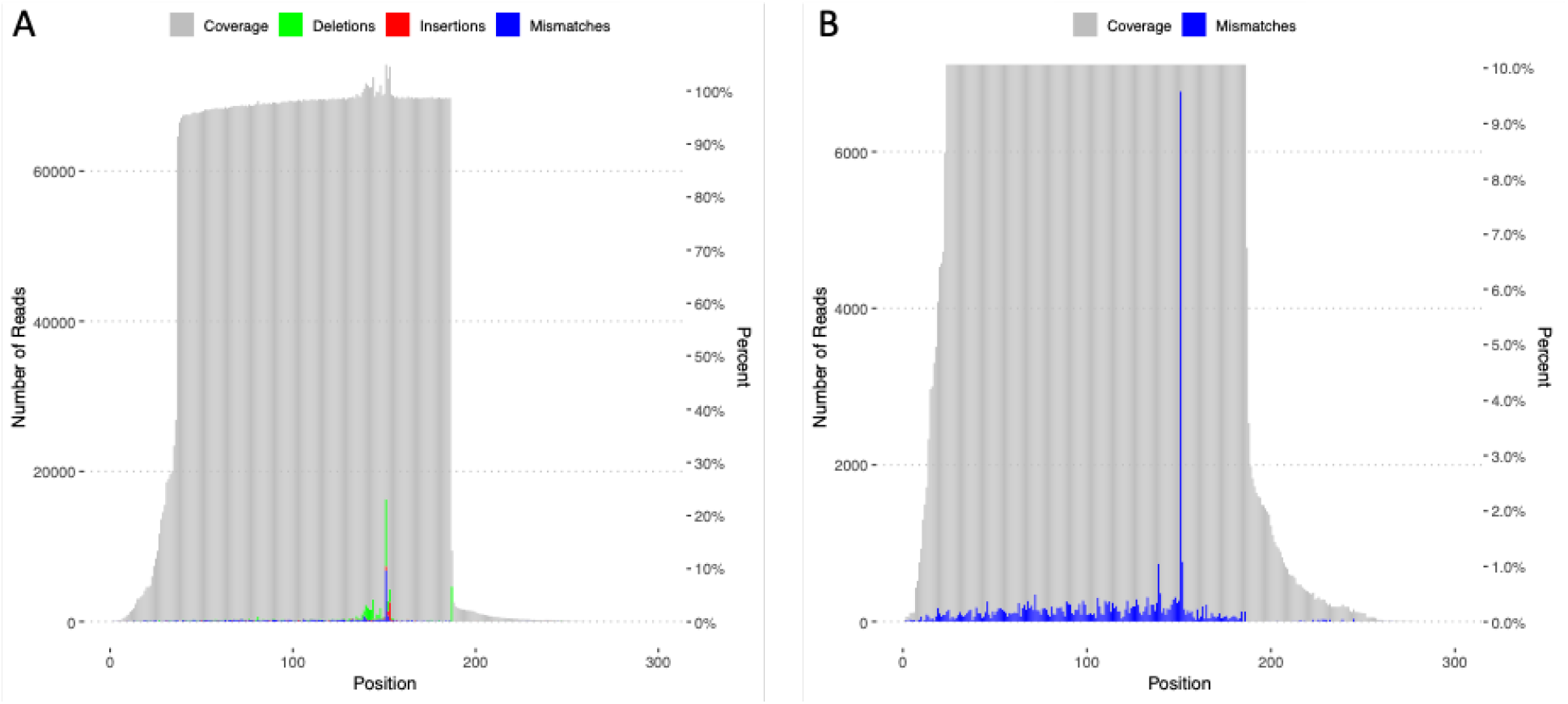
An example of EdiTyper summary plots for real sequencing data from a polyclonal cell population from CRISPR experiment. A) The main locus landscape plot summarizing all editing events and sequencing coverage. B) Landscape plot showing only mismatches, with the high peak corresponding to the SNP introduced by homologous recombination. Analogous plots are provided also for insertions and deletions (data not shown).

## 4 Discussion

In this paper we presented EdiTyper, an open source software tool that characterizes targeted sequencing data from genome editing experiments with highly robust and accurate results and high processing speed. EdiTyper is robust to sequencing noise from different Illumina MiSeq versions, with high accuracy even for high-error platforms like MSv3. EdiTyper compares favorably to CRISPResso, a previously published tool for similar purposes, with better or equal performance, sometimes with dramatic improvements. EdiTyper is also an extremely fast pipeline, benefitting from a very fast custom aligner well integrated in a light Python software. This intrinsic speed is particularity relevant for the increasing scope and scale of CRISPR genome editing experiments. EdiTyper is mainly designed and tested for CRISPR experiments, but should be applicable to other genome editing approaches as well. The open-source software is built to minimize dependencies to enable easy installation and maintenance, is parallelizable, and the flexible command-line interface includes many adjustable parameters and options. Altogether, EdiTyper offers highly accurate analysis of sequencing data from genome editing experiments, and is applicable to large volumes of experimental data from increasingly large and diverse genome editing experiments.

## Acknowledgments

We would like to thank Ana Vasileva, Sarah Kim-Hellmuth, Stephane Castel, and other members of the Lappalainen Lab for testing EdiTyper and providing helpful comments and suggestions. T.L. was supported by the NIH grants R01GM122924, R01MH106842, UM1HG008901 and 1U24DK112331. P.M. was supported by the NIMH grant R01MH106842. A.Y. was supported by the NIGMS grant R01GM107145. N.P.T. was supported by the NIGMS grant R01GM107145 and the NCATS grant OT3TR002027. P.M. was supported by CTSA grant UL1TR002550. T.L. is a member of the scientific advisory board of Goldfinch Bio and Variant Bio, and has equity in Variant Bio.

