## Supplementary material for "EdiTyper: a high-throughput tool for analysis of targeted sequencing data from genome editing experiments"

### A Supplementary Material

#### A.1 Supplementary tables

**Table S1.** 17 reference protein-coding gene loci of 150bp used for the benchmarking experiments. Available in .csv format attached to this paper.

| | **No error** | | **MSv1** | | **MSv3** | |
|---|---|---|---|---|---|---|
| | precision | recall | precision | recall | precision | recall |
| HDR | 0.8738 (±0.2152) | 1.0 (±0.0) | 0.8484 (±0.2087) | 0.9974 (±0.0059) | 0.9275 (±0.0973) | 0.4397 (±0.0575) |
| NHEJ | 0.9996 (±0.0005) | 0.8673 (±0.064) | 0.7952 (±0.0155) | 0.8938 (±0.051) | 0.4923 (±0.0175) | 0.987 (±0.0044) |
| MIX | 0.75 (±0.4331) | 0.7442 (±0.4299) | 0.9505 (±0.141) | 0.7527 (±0.4119) | 0.6138 (±0.0388) | 0.9697 (±0.0432) |
| UNC | 0.8959 (±0.0482) | 1.0 (±0.0) | 0.8958 (±0.0484) | 0.794 (±0.0066) | 0.8888 (±0.0382) | 0.0865 (±0.0046) |

**Table S2.** Average precision and recall (with standard deviation) for HDR, NHEJ, MIX (i.e., HDR with indel) and UNC (unchanged reads) for CRISPResso across error profiles.

| | **No error** | | **MSv1** | | **MSv3** | |
|---|---|---|---|---|---|---|
| | precision | recall | precision | recall | precision | recall |
| HDR | 0.9799 (±0.0172) | 1.0 (±0.0) | 0.9488 (±0.0497) | 0.9977 (±0.0055) | 0.9488 (±0.0495) | 0.9739 (±0.0177) |
| NHEJ | 0.9993 (±0.0009) | 0.9785 (±0.0073) | 0.9991 (±0.001) | 0.9778 (±0.0073) | 0.972 (±0.0041) | 0.9779 (±0.0069) |
| MIX | 0.9768 (±0.0544) | 0.9681 (±0.0249) | 0.949 (±0.0778) | 0.9659 (±0.0254) | 0.9259 (±0.0781) | 0.9659 (±0.0247) |
| UNC | 0.9815 (±0.0064) | 1.0 (±0.0) | 0.9815 (±0.0063) | 0.9992 (±0.0005) | 0.9811 (±0.0061) | 0.9747 (±0.0029) |

**Table S3.** Average precision and recall (with standard deviation) for HDR, NHEJ, MIX (i.e., HDR with indel) and UNC (unchanged reads) for EdiTyper across error profiles.

##### A.2 Supplementary figures

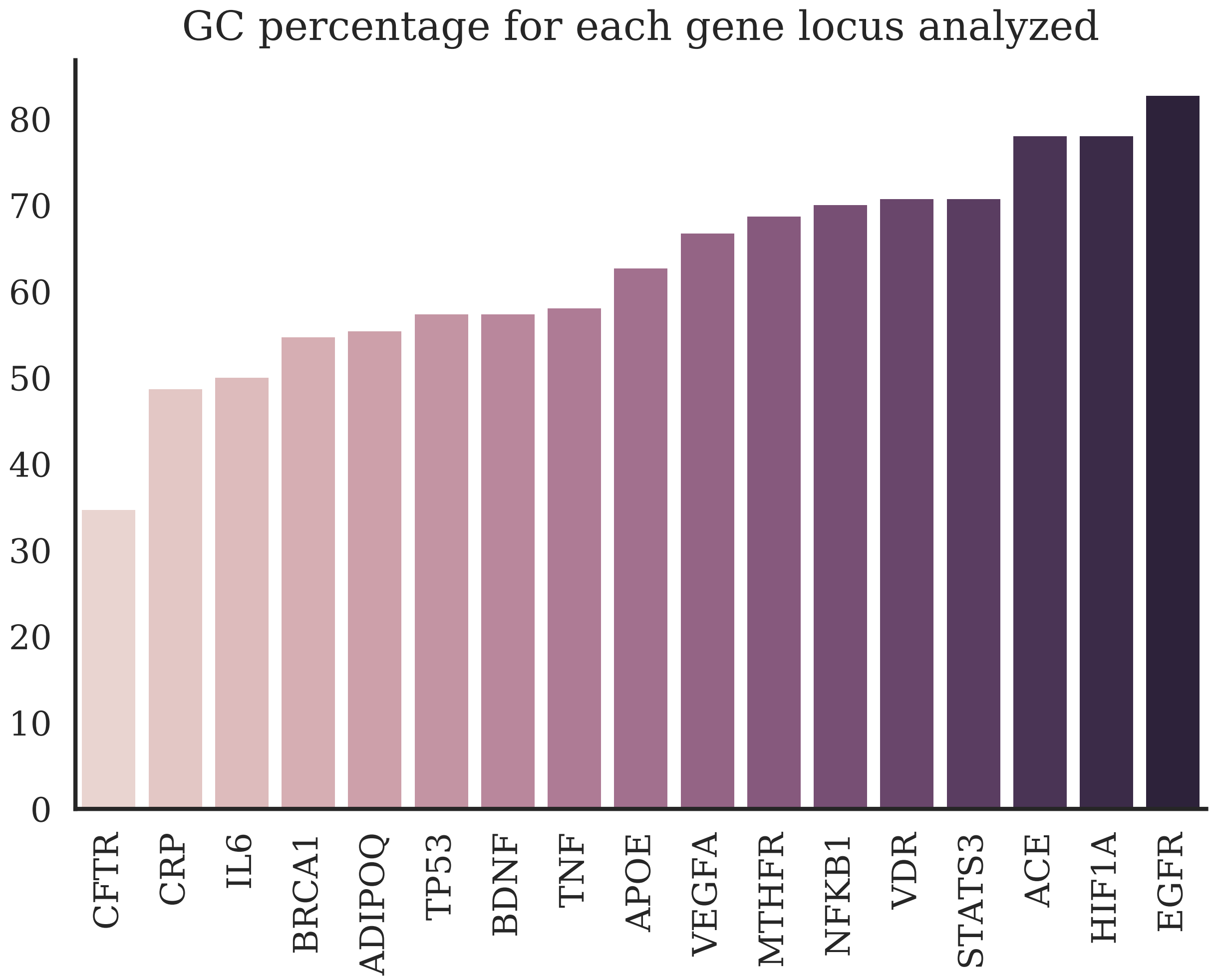


**Figure S1.** GC% of each protein-coding gene locus selected for our experiments

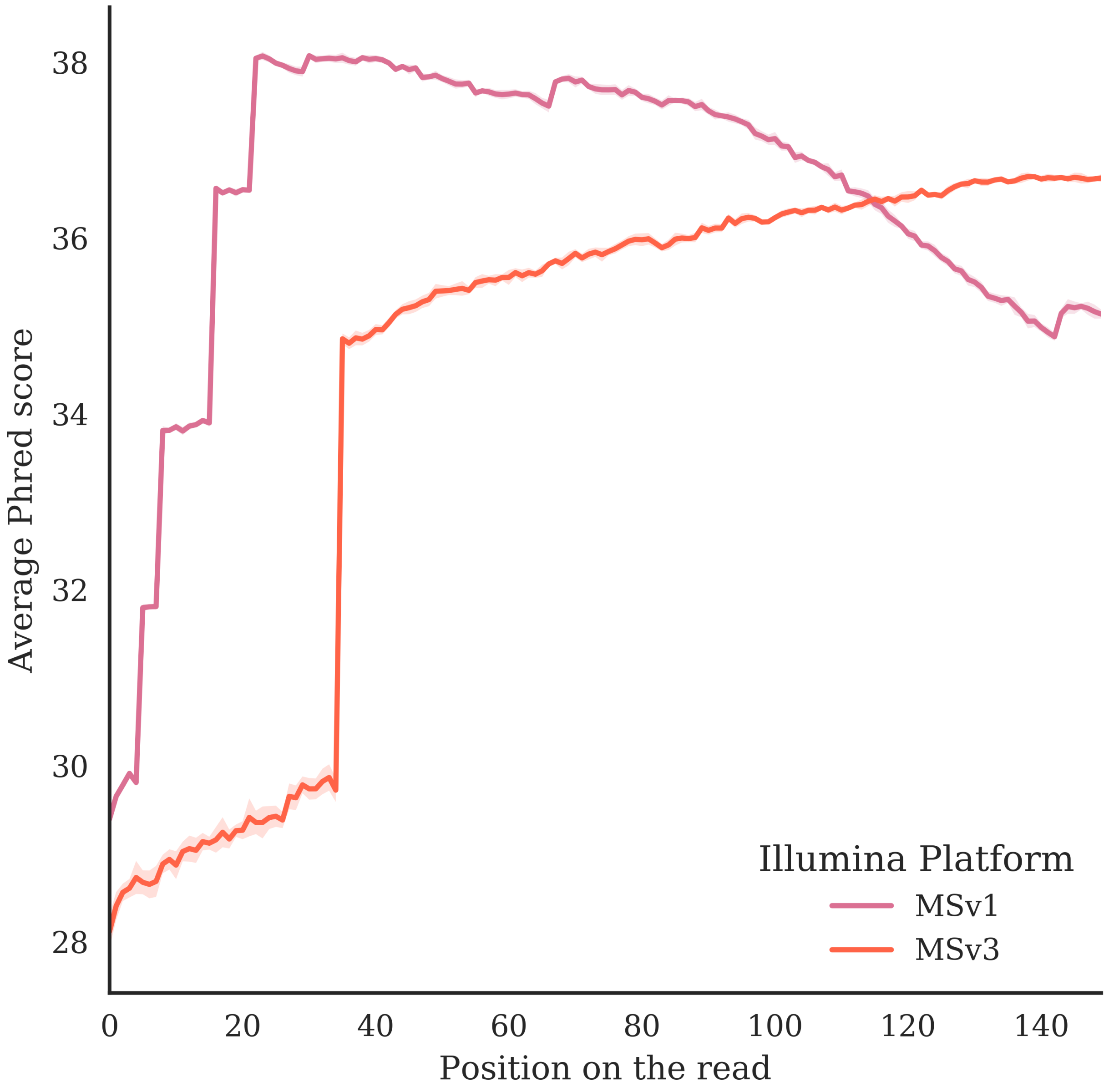


**Figure S2.** Average Phred score by position on the read for generated sequencing datasets with the error profiles of two Illumina platforms simulated with ART.

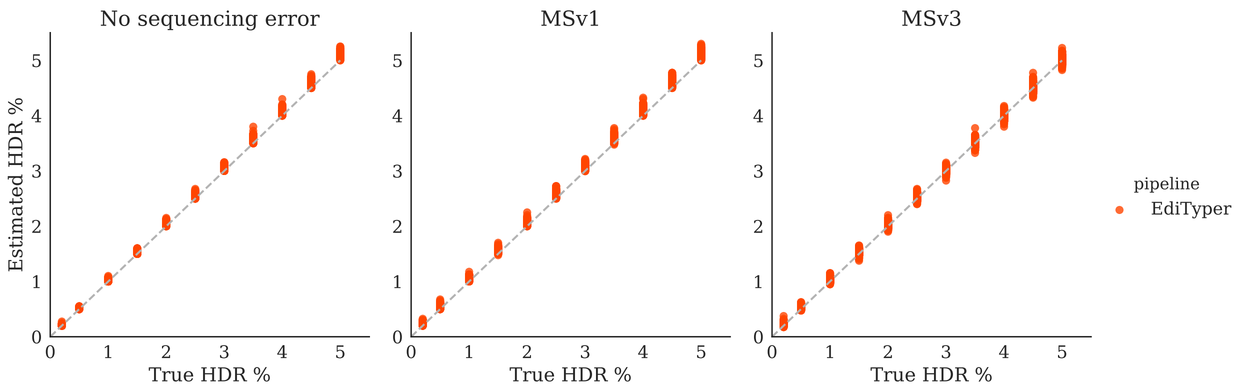


**Figure S3.** True HDR percentage vs. estimated percentage by EdiTyper for datasets without sequencing error profile, MSv1 error profile, and MSv3 error profile.

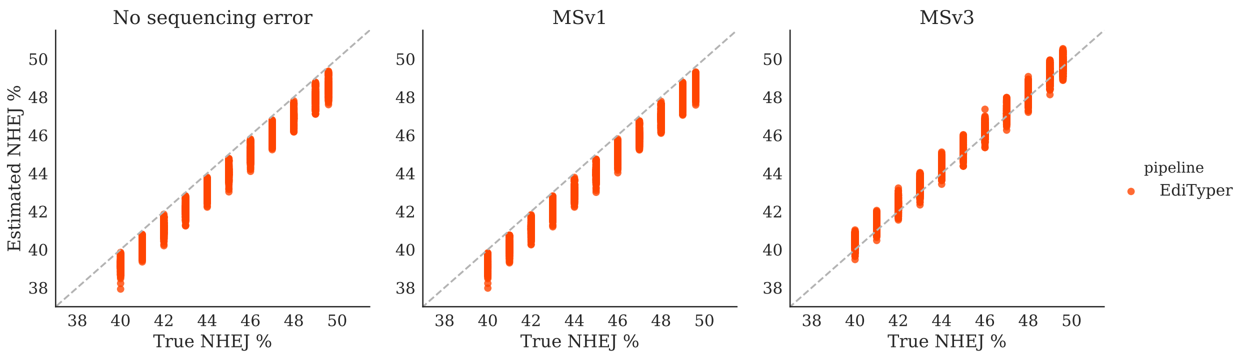


**Figure S4.** True NHEJ percentage vs. estimated percentage by EdiTyper for datasets without sequencing error profile, MSv1 error profile, and MSv3 error profile.

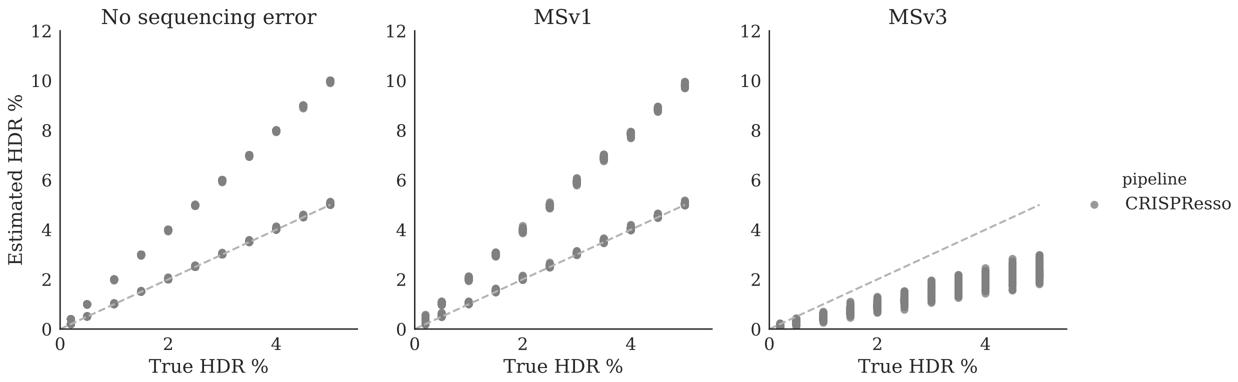


**Figure S5.** True HDR percentage vs. estimated percentage by CRISPResso for datasets without sequencing error profile, MSv1 error profile, and MSv3 error profile. Over-estimated HDR percentages were from datasets with 1 bp indel.

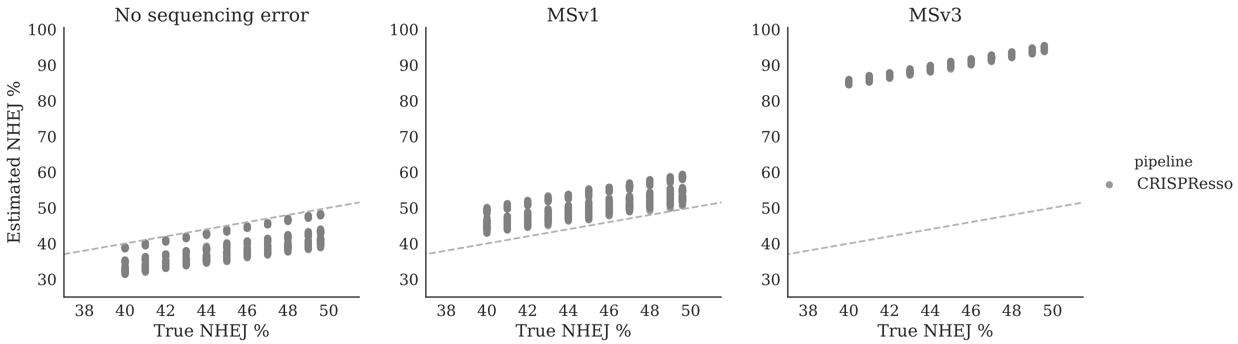


**Figure S6.** True NHEJ percentage vs. estimated percentage by CRISPResso for datasets without sequencing error profile, MSv1 error profile, and MSv3 error profile.

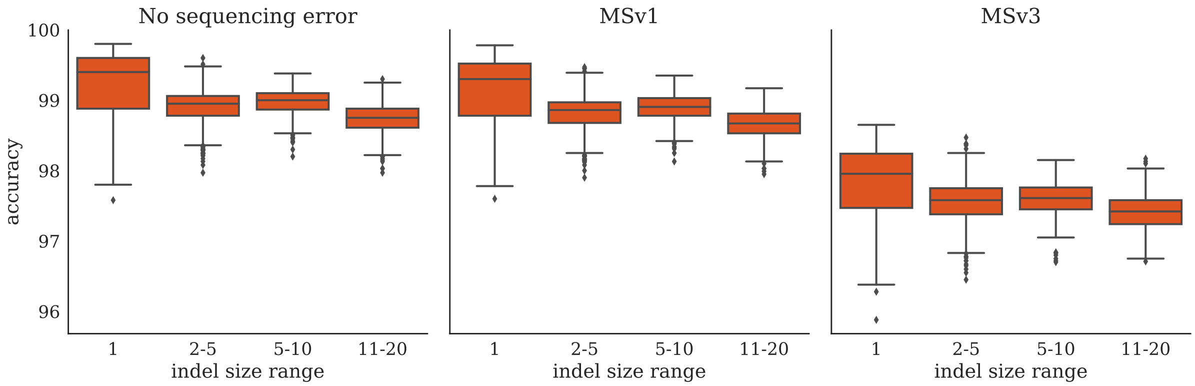


**Figure S7.** EdiTyper's accuracy in function of the average indel size for each sequencing error profile. Accuracy was negatively correlated with average indel size (Pearson $r = -0.19$, p-value$< 10e - 69$).

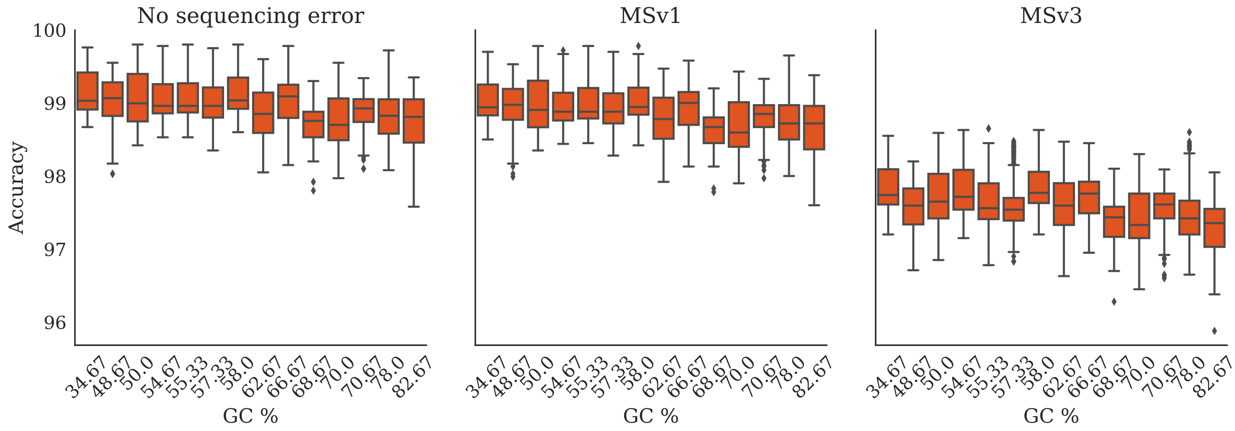


**Figure S8.** EdiTyper's accuracy in function of the GC percentage of the seed locus used to generate the datasets. Accuracy was overall negatively correlated with the GC percentage of the seed locus (Pearson $r = -0.78$, p-value$= 0.0002$).

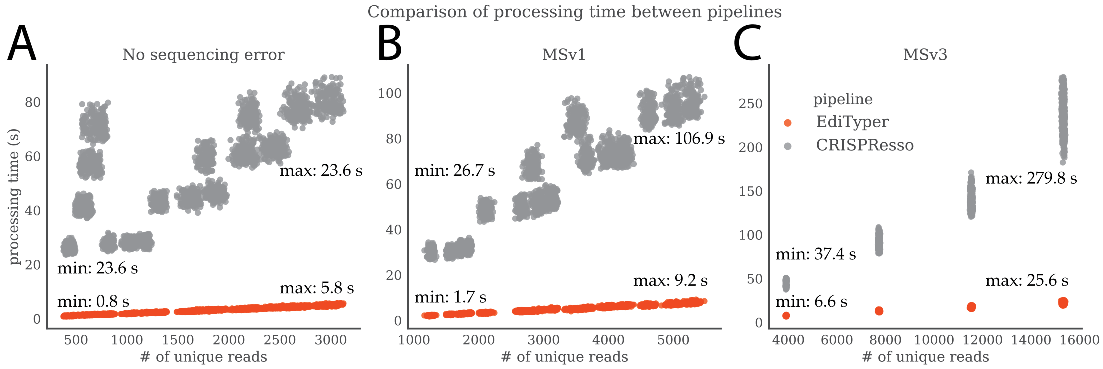


**Figure S9.** Comparison of the speed performances between EdiTyper and CRISPResso. (A) Processing time as a function of the number of unique reads in each dataset with no sequencing error, (B) MSv1 error profile, and (C) MSv3 error profile. Sequencing errors increase the number of unique reads.
